# Genetically engineered orange petunias on the market

**DOI:** 10.1101/142810

**Authors:** Hany Bashandy, Teemu H. Teeri

## Abstract

Genetic engineering of petunia was shown to lead to novel flower color some twenty years ago. Here we show that petunia lines with orange flowers, generated for scientific purposes, apparently found their way to petunia breeding programmes, intentionally or unintentionally. Today they are widely available, but have not been registered for commerce.

---

The pathway to the colored anthocyanins in the ornamental plant petunia *(Petunia hybrida)* is a well-known example of substrate specificity of one enzyme limiting the spectrum of possible products of the pathway^1^. Anthocyanins are water soluble pigments giving flowers, fruits and sometimes vegetative parts of plants colours ranging from orange and red to blue and purple^2^. Anthocyanins are extensively glycosylated and acylated, the molecular decoration affecting their spectral properties. At the aglycone level the three most common variants of the molecule are the anthocyanidins pelargonidin, cyanidin and delphinidin, differing by the number of hydroxyl groups (one, two or three, respectively) in the B-ring of the molecule. Hydroxylation takes place at the level of dihydroflavonols in the pathway (possibly earlier in some cases) by two enzymes, flavonoid 3’-hydroxylase (F3’H) and flavonoid 3’5’-hydroxylase (F3’5’H). The enzyme dihydroflavonol reductase (DFR) converts dihydroflavonols to corresponding leucoanthocyanidins, which then are oxidized to anthocyanidins by anthocyanidin synthase (syn. leucoanthocyanidin oxidase). In petunia, the DFR enzyme does not react with the simplest precursor (dihydrokaempferol), therefore the natural range of petunia flower colours lack orange hues typical to pelargonidin derivatives. Flowers of petunia cultivars that have mutations in the two hydroxylases are therefore white.

It was shown few decades ago that by introducing a gene encoding DFR from a species where the enzyme does not show substrate specificity into a petunia line that lacks F3’H and F3’5’H activity, one can open up the pathway to pelargonidin. Using the maize gene *A1* Meyer and colleagues generated brick red colored flowers in petunia^3^ and using the gene from the ornamental plant *Gerbera hybrida,* our own laboratory generated petunia lines with bright orange flowers^4^.

These petunia flowers were investigated concerning factors relating to stability of the transgene (and therefore the novel colour)^5,6^, but they were never commercialized. The list of registered genetically modified petunia plants is very short and includes a single line transgenic for a chalcone synthase encoding gene approved for cultivation in China (http://www.isaaa.org/gmapprovaldatabase/). Therefore, it was a great surprise and a delight from the point of view of maybe gaining insight in the ways petunia germplasm changes under breeding, when we encountered bright orange coloured petunias in flower boxes decorating the Helsinki railway station during summers of 2015 and 2016 (Figure 1). Indeed, orange petunias are widely on the market, as an internet search with these keywords shows. The cultivar at the Helsinki railways station was “Bonnie Orange”, for which we got a sample from the gardeners of the City of Helsinki in 2016. The cultivar “African Sunset” was available from several sources, along with a number of others that we were able to purchase (Table 1 and Figure S1).

**Fig. 1.**
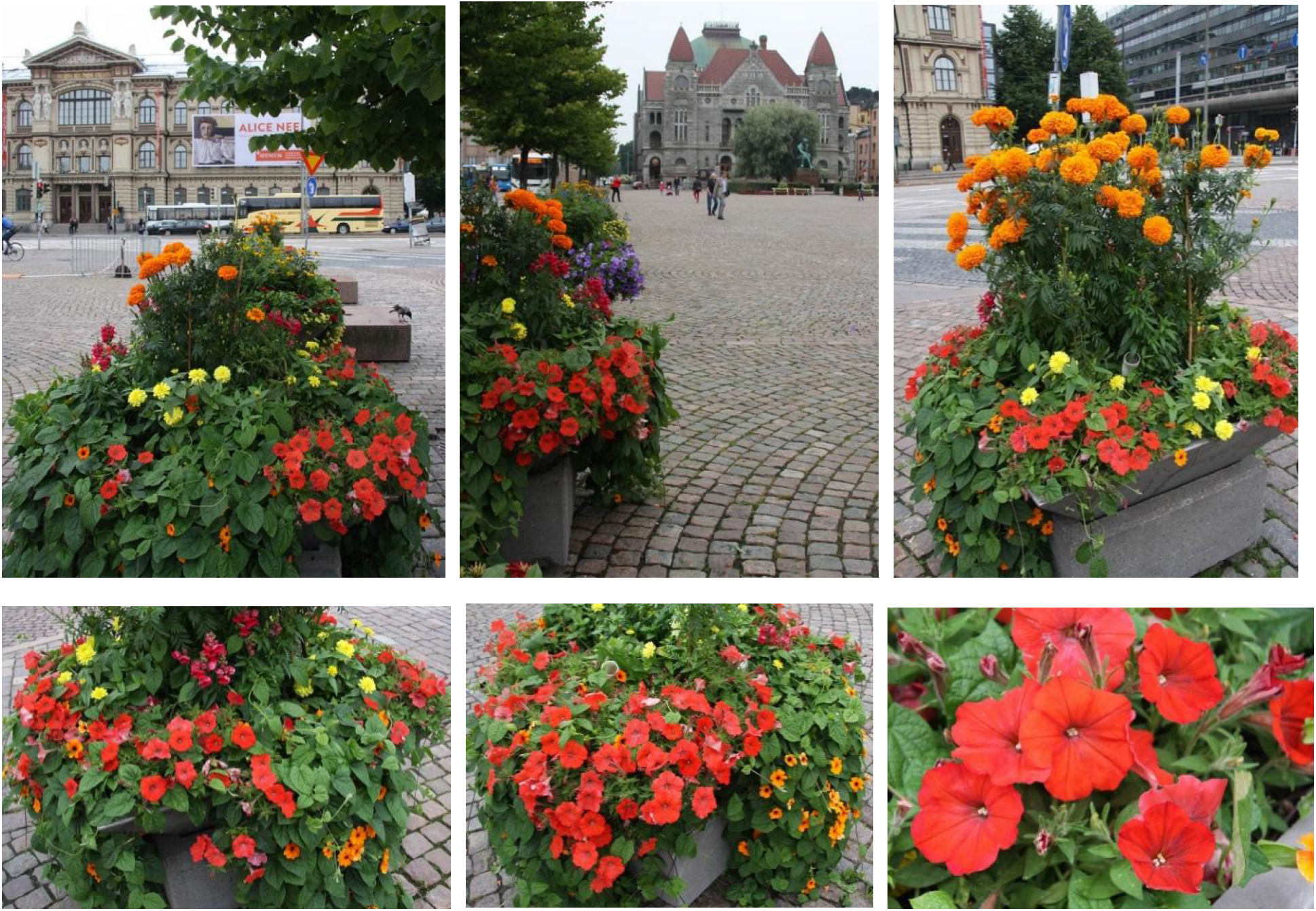
Orange petunias decorating the Helsinki railway station in 2016.

**Table 1.** “Orange colored” petunias purchased for this study

| Table 1. “Orange colored” petunias purchased for this study |  |  |  |  |
| --- | --- | --- | --- | --- |
| Number | Cultivar name | Source | Main anthocyanidin | Comments |
| 1 | Cascadias Indian Summer | City of Helsinki | Cyanidin | Low amounts of anthocyanins |
| 2 | Bonnie Orange | City of Helsinki | Pelargonidin |  |
| 3 | African Sunset | Amazon.com<sup>1</sup> | Pelargonidin |  |
| 4 | African Sunset | Nicky’s Nursery<sup>2</sup> | Pelargonidin |  |
| 5 | African Sunset | Etsy<sup>3</sup> |  | Did not germinate |
| 6 | African Sunset | Etsy<sup>3</sup> | Pelargonidin |  |
| 7 | African Sunset | Swallowtail Garden<sup>4</sup> | Pelargonidin |  |
| 8 | Alladdin Orange | Seedman<sup>5</sup> | Cyanidin |  |
| 9 | Orange color Petunia | Aliexpress<sup>6</sup> |  | Flowers purple or white with purple stripes |
| 10 | Rare orange petunia | Aliexpress<sup>6</sup> |  | Did not germinate |
| Web sites for seed sources |  |  |  |  |
| <sup>1</sup>https://www.amazon.com/African-Sunset-Beautiful-Orange-Petunia/dp/B00TG0E3PI |  |  |  |  |
| <sup>2</sup>http://www.nickys-nursery.co.uk/garden-shop/seeds/hanging-baskets/petunia/petunia.-african-sunset-10-pellets |  |  |  |  |
| <sup>3</sup>https://www.etsy.com/cart/ |  |  |  |  |
| <sup>4</sup>http://www.swallowtailgardenseeds.com/annuals/petunia.html#gsc.tab=0 |  |  |  |  |
| <sup>5</sup>https://www.seedman.com/petunia.htm |  |  |  |  |
| <sup>6</sup>http://www.aliexpress.com/price/orange-petunia-seeds_price.html |  |  |  |  |

Chemical analysis showed that while “Aladdin Orange” with orange red flowers was a cyanidin cultivar, “Bonnie Orange” and “African Sunset” unexpectedly contained pelargonidin as the main anthocyanidin. (Table 1, Figure S2). Based on what we knew about petunia flower colour and the history of orange petunias, we made a quick test using primers designed for the maize *A1* gene (using a method called reverse transcription PCR) and saw that the orange flowers of “Bonnie Orange” and “African Sunset” indeed expressed a maize gene – and in addition the most common gene transfer selective marker gene *nptII* (Fig S3). We further made a guess that these transgenic lines might contain the widely used CaMV 35S promoter sequence driving either of the two transgenes, and perhaps also the *bla* gene (for ampicillin resistance in bacterial hosts) common in many vectors. We designed primers that would amplify these sequences, as well as primers that would initiate amplification out of these sequences to nearby genetic elements (Table S1). Using an array of petunia chromosomal DNA samples (lines 1-4 and 6-9 in Table 1) and different combinations of the primers, we could amplify not only the *A1* and *nptII* genes, but also sequences between 35S and A1, *A1* and *nptII,* and *bla* and 35S. Amplification of these fragments took place repeatedly and exclusively from chromosomal DNA extracted from leaves of “Bonnie Orange” and “African Sunset” (Fig S4). The PCR fragments were directly sequenced and due to presence of overlapping parts they could be assembled into a single contig corresponding very precisely to the map presented in Meyer et al.^3^. We could not amplify the *bla* gene as a whole or sequences between the *bla* gene and the *nptII* gene, a possible explanation being that this region represents the integration site of the plasmid in a petunia chromosome. In fact, the line described in detail by Meyer and colleagues^5^ and used by Oud and colleagues in breeding experiments (see below)^7^ with a single copy insert and a truncated *bla* gene fits with these observations.

The retrieved sequences were exactly identical in “Bonnie Orange” and “African Sunset”, indicating a common origin. Phenotypically these two cultivars are not exactly alike, for example “African Sunset” has larger flowers than “Bonnie Orange”. The recipient line Meyer and colleagues^3^ used in their experiment was chosen based on its mutant genotype lacking both of the two hydroxylases (*ht ht hf hf*). It is not a line with good horticultural properties, but as with other interesting characters with simple inheritance, petunia breeders could introgress the gene for orange petal color to horticulturally superior genetic background by simple crossing. This was successfully done by Oud and colleagues^7^, showing that trait improvement by traditional breeding indeed works well also for genetically modified traits. The article ends by stating that “[the orange colour trait] has been successfully used in breeding programmes aimed at developing commercial F1 varieties with this trait”. This, relying on records, never happened. The regulations in Europe and elsewhere require an extensive analysis of risks a genetically modified organism might impose on human health and the environment upon deliberate release (i.e., commercialization). This expensive procedure prohibits the use of genetic modification in cases where the expected volume of production would be too small to cover the extra expenses – obviously the case for a petunia cultivar with a novel colour.

The orange petunia lines generated for scientific purposes apparently found their way to petunia breeding programmes, intentionally or unintentionally. This particular escaped genetically engineered (GM) line causes no harm, but demonstrates that containment procedures can never be made failproof. A more important observation is that regulation of GM crops based on the breeding method instead of the cultivar’s properties in practise completely prohibits commercialisation of lines with traits that have beneficial but only incremental value – the typical pattern of gain in plant breeding. Although orange coloured petunias are mere examples of breeding for beauty, for developing better crops the lost opportunities of GM breeding have much wider consequences.

## Methods

### Chemicals and plant material

Authentic standards for pelargonidin and cyanidin were purchased from TransMIT PlantMetaChem (Giessen, Germany). All other chemicals were from Sigma-Aldrich (Milano, Italy), water was purified by a Milli-Q water purification system. Petunia seeds were germinated and plants were grown in peat:vermiculite (1:1) under fluorescent illumination (16 h day length) at 23 °C. Origin of the different petunia cultivars is shown in Table 1, the nontransgenic line W80^8^ was a kind gift of Dr. Ronald Koes.

### RNA extraction, cDNA synthesis and RT-PCR reactions

Total RNA was isolated as described^9^ from unopened flowers when anthocyanin biosynthesis was active and the petals were gaining color. A treatment with RNase free DNase (Nucleo Spin RNA clean-up XS, Macherey-Nagel, Germany) was done to remove any residues of genomic DNA. First-strand cDNA was synthesized from 166 ng of total RNA using the Superscript III Reverse Transcriptase Kit (Invitrogen). Using the cDNA as template, the full length maize *A1* transcript was amplified with primers GER945 and GER946, and a fragment of the *nptII* transcript with primers GER522 and GER523 using Phusion High-Fidelity DNA Polymerase (Thermo Scientific, Waltham, MA, USA). The thermal cycler was programmed in the following way: An initial cycle of denaturation at 98 °C for 30 s, followed by 25 (A1) or 30 *(nptII)* cycles of: 98 °C for 30 s, 65°C (A1) or 56°C *(nptII)* for 30 s, 72°C for 1 min, followed by a final extension for 10 min at 72 °C.

### DNA extraction and genomic PCR

Genomic DNA was isolated from leaves using the miniprep II method^10^. (1983). 100 ng of petunia genomic DNA was used for amplification of sequences between the *bla* gene and the 35S promoter (primers GER1003 and GER992), between the 35S promoter and the maize *A1* gene (primers GER993 and GER984) and between the maize *A1* gene and the *nptII* gene (primers GER981 and GER976) with Dream Taq DNA polymerase (Thermo Scientific, Waltham, MA, USA) with the following program: An initial cycle of denaturation at 95 °C for 3 min, followed by 30 cycles of: 94 °C for 1 min, 55°C for 30 s, 72°C for 2 min. Amplified fragments were purified using High Pure PCR Product Purification Kit (Roche Diagnostics, Indianapolis, IN, USA) and directly sequenced using the amplification primers and, when needed, internal primers designed from the sequences. The sequences were assembled together based on their overlapping segments and deposited in GenBank with the accession number KY964325.

### Anthocyanidin analysis

HPLC analysis was carried out as described^11^ with minor modifications. Samples were collected from fully opened petunia flowers, ground in liquid nitrogen and extracted with a 2x volume of methanol with 1% HCl. After sonication for 30 min at room temperature (Finn sonic W181, Oy ULTRA sonic Finland Ltd, Lahti, Finland), the extracts were cleared from debris by centrifugation (3220 ×g, 10 min). The supernatant was mixed 1:1 with 4 M HCl and hydrolyzed by incubating for 40 min at 95°C. The hydrolyzed extracts were centrifuged 17000 ×g for 10 min before injection to HPLC. The standard compounds were applied at 10 μg/ml in methanol.

## Supplementary information

**Fig. S1.**
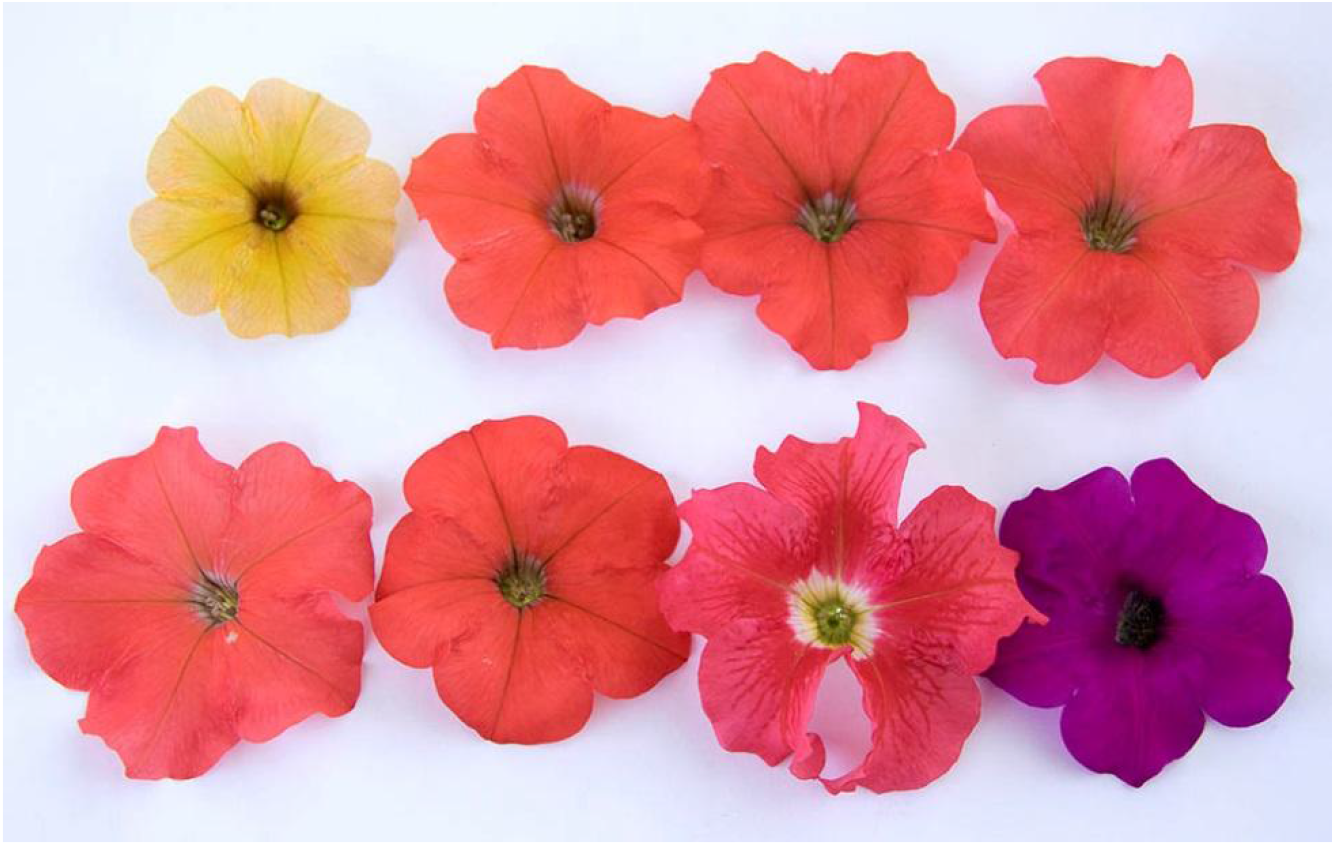
Petunia cultivars “Indian Summer”, “Bonnie Orange”, “African Sunset” (4 sources), “Aladdin Orange” and “Orange color petunia”.

**Fig. S2.**
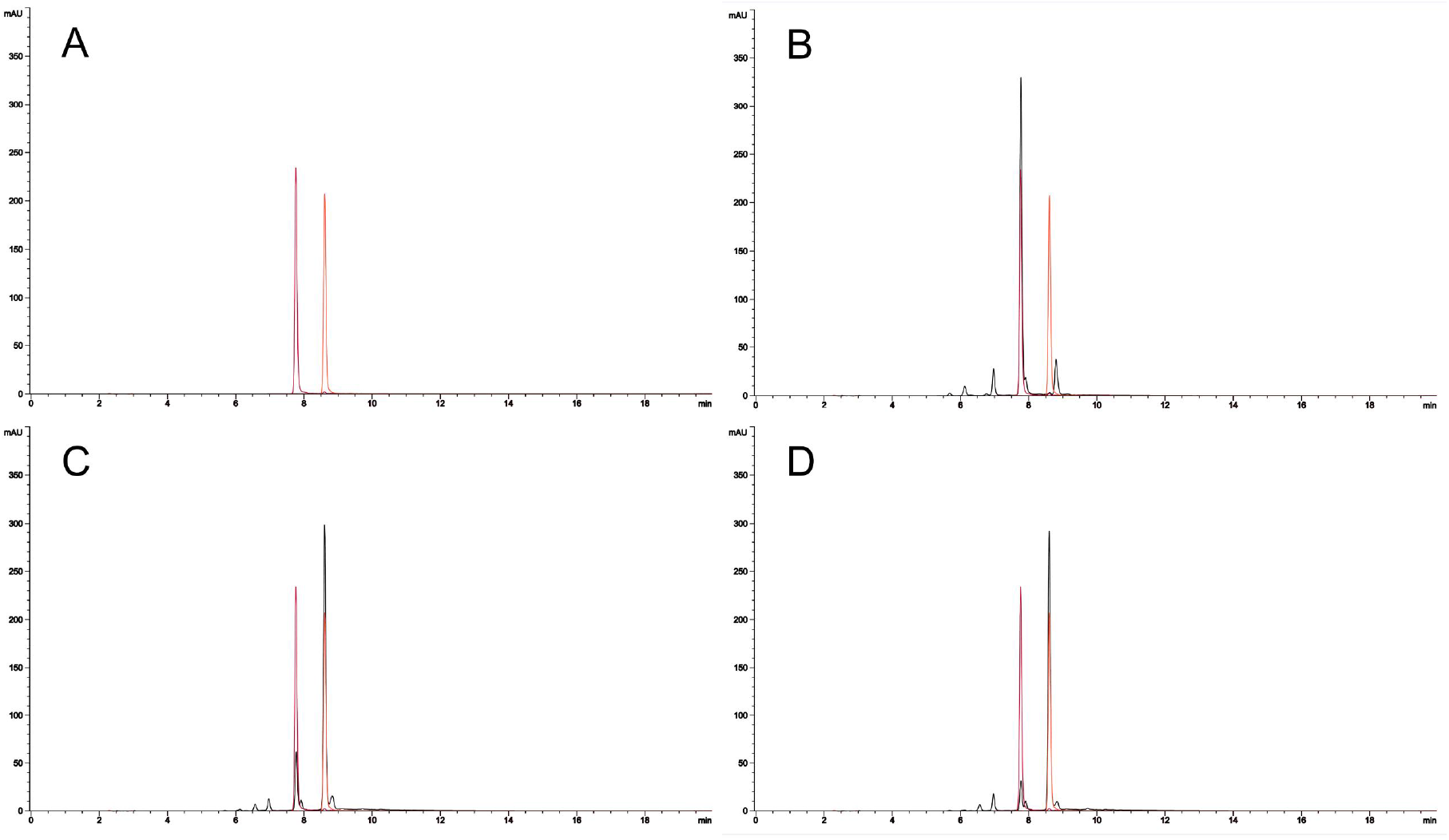
HPLC chromatograms of anthocyanidins extracted from orange petunia flowers. (**A**). Authentic standards for cyanidin (left, 7.80 min) and pelargonidin (right, 8.64 min). (**B**). Cultivar “Aladdin Orange”, containing cyanidin derived anthocyanidins. (**C**). Cultivar “Bonnie Orange”, containing pelargonidin derived anthocyanidins. (**D**). Cultivar “African Sunset”, containing pelargonidin derived anthocyanidins. Panels B to D are overlayed with the chromatograms of the authentic standards (in color).

**Fig. S3.**
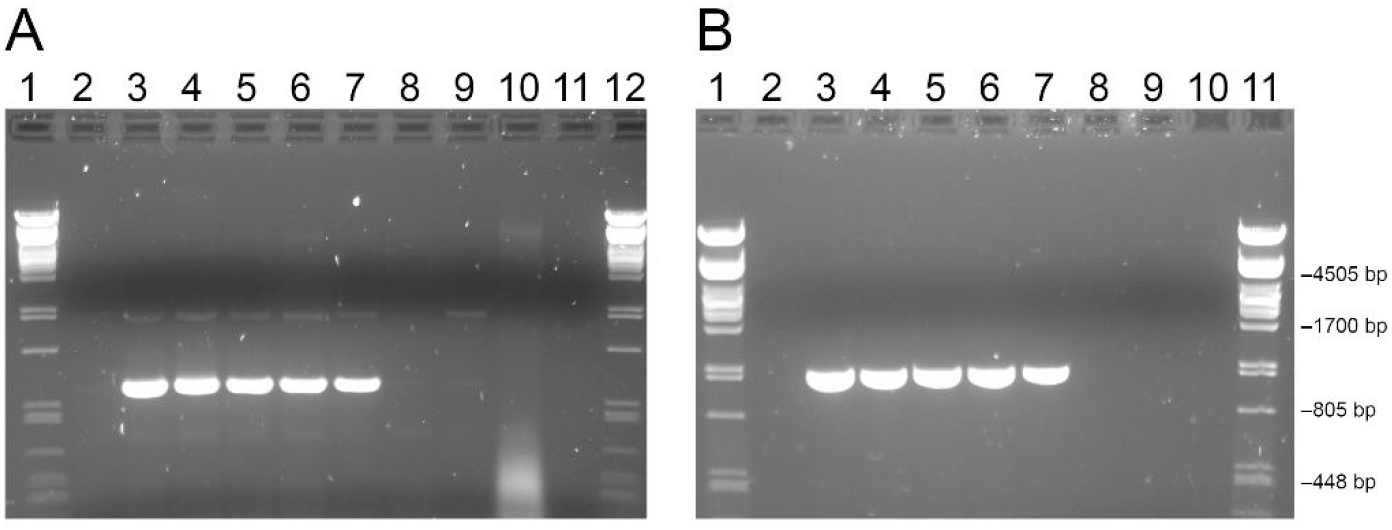
RT-PCR from petunia mRNA. Outermost lanes: PstI digested Lambda DNA. (**A**). Primers amplifying *nptII* sequences. Lanes 2-11: “Cascadias Indian Summer”, “Bonnie Orange”, “African Sunset” (four sources), “Alladin Orange”, “W80”, “Orange color Petunia”, water. (**B**). Primers amplifying the maize *A1* DFR sequences. Lanes 2-10: “Cascadias Indian Summer”, “Bonnie Orange”, “African Sunset” (four sources), “Alladin Orange”, “W80”, water.

**Fig. S4.**
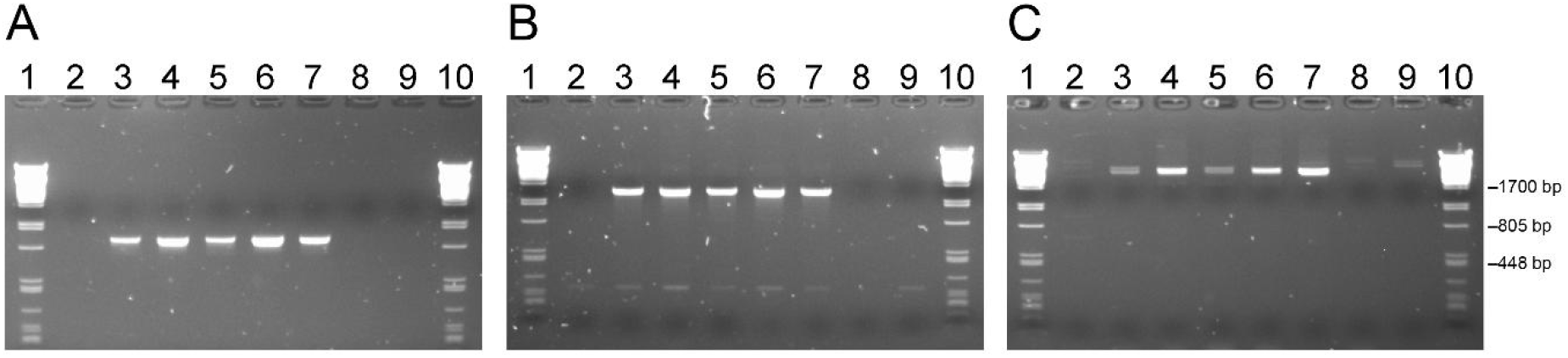
PCR from petunia genomic DNA. Outermost lanes: PstI digested Lambda DNA. Lanes 2-9: “Cascadias Indian Summer”, “Bonnie Orange”, “African Sunset” (four sources), “Alladin Orange”, “Orange color Petunia”. (**A**). Primers amplifying sequences from *bla* to the 35S promoter. (**B**). Primers amplifying sequences from the 35S promoter to the maize *A1* DFR. (**C**). Primers amplifying sequences from the maize *A1* DFR to *nptII*.

**Table S1.**
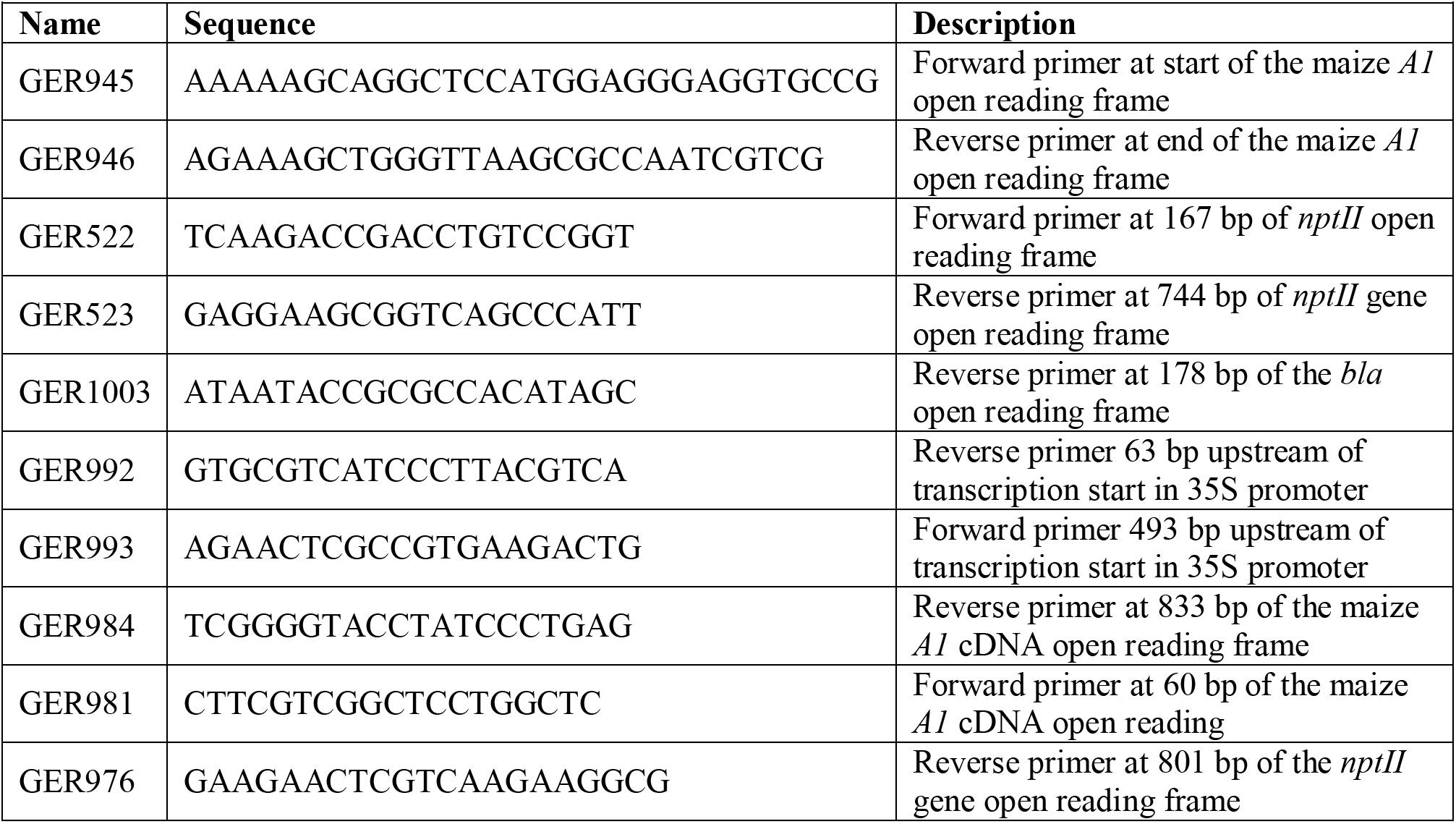
Primers used in this study

## References

1. G. Forkmann, G. & Ruhnau, B. Distinct substrate specificity of dihydroflavonol 4-reductase from flowers of Petunia hybrida. Zeitschrift für Naturforschung C 42, 1146–1148 (1987).

2. Winkel-Shirley, B. Flavonoid biosynthesis. A colorful model for genetics, biochemistry, cell biology, and biotechnology. Plant Phys. 126, 485–493 (2001).

3. Meyer, P., Heidmann, I., Forkmann, G. & Saedler, H. A new petunia flower colour generated by transformation of a mutant with a maize gene. Nature 330, 677–678 (1987).

4. Helariutta, Y., Elomaa, P., Kotilainen, M., Seppänen, P. & Teeri, T.H. Cloning of cDNA coding for dihydroflavonol-4-reductase (DFR) and characterization of dfr expression in the corollas of Gerbera hybrida var. Regina (Compositae). Plant Mol. Biol. 22, 183–193 (1993).

5. Meyer, P., Linn, F., Heidmann, I., Meyer, H., Niedenhof, I. & Saedler, H. Endogenous and environmental factors influence 35S promoter methylation of a maize *A1* gene construct in transgenic petunia and its colour phenotype. Mol. Gen. Gen. 231, 345–352 (1992).

6. Elomaa, P., Helariutta, Y., Kotilainen, M., Teeri, T.H., Griesbach, R.J. & Seppänen, P. Transgene inactivation in Petunia hybrida is influenced by the properties of the foreign gene. Mol. Gen. Gen. 248, 649–656 (1995).

7. Oud, J.S., Schneiders, H., Kool, A.J. & van Grinsven, M.Q. Breeding of transgenic orange Petunia hybrida varieties. Euphytica 84, 175–181 (1995).

8. Huits, H.S., Gerats, A.G., Kreike, M.M., Mol, J.N. & Koes, R.E. Genetic control of dihydroflavonol 4-reductase gene expression in Petunia hybrida. Plant J. 6, 295–310 (1994).

9. Chang, S., Puryear, J. & Cairney, J. A simple and efficient method for isolating RNA from pine trees. Plant Mol. Biol. Rep. 11, 113–116 (1993).

10. Dellaporta, S.L., Wood, J. & Hicks, J.B. A plant DNA minipreparation: version II. Plant Mol. Biol. Rep. 1, 19–21 (1983).

11. Bashandy, H., Pietiäinen, M., Carvalho, E., Lim, K.J., Elomaa, P., Martens, S. & Teeri, T.H. Anthocyanin biosynthesis in gerbera cultivar ‘Estelle’and its acyanic sport ‘Ivory’. Planta 242, 601–611 (2015).

